# Improvement of glymphatic-lymphatic drainage of beta-amyloid by focused ultrasound in Alzheimer’s disease model

**DOI:** 10.1101/2020.01.24.918607

**Authors:** Youngsun Lee, Yoori Choi, Eun-Joo Park, Seokjun Kwon, Hyun Kim, Jae Young Lee, Dong Soo Lee

## Abstract

Drainage of parenchymal waste through the lymphatic system maintains brain homeostasis. Age-related changes of glymphatic-lymphatic clearance lead to the accumulation beta-amyloid (Aβ) in dementia models. In this study, focused ultrasound treatment in combination with microbubbles (FUS-MB) improved Aβ drainage in early dementia model mice, 5XFAD. FUS-MB enhanced solute Aβ clearance from brain, but not plaques, to cerebrospinal fluid (CSF) space and then deep cervical lymph node (dCLN). dCLN ligation exaggerated memory impairment and progress of plaque formation and also the beneficial effects of FUS-MB upon Aβ removal through CSF-lymphatic routes. In this ligation model, FUS-MB improved memory despite accumulation of Aβ in CSF. In conclusion, FUS-MB enhances glymphatic-lymphatic clearance of Aβ mainly by increasing brain-to-CSF Aβ drainage. We suggest that FUS-MB can delay dementia progress in early period and benefits of FUS-MB depend on the effect of Aβ disposal through CSF-lymphatics.

## Introduction

The glymphatic-lymphatic system plays an important role in the clearance of extracellular metabolites and waste products in the brain^1, 2^. The glymphatic (glial-lymphatic) is a postulated system for the CSF-interstitial fluid (ISF) exchange in the brain driven by the CSF influx force, which moves solutes from the periarterial CSF space via ISF efflux to the perivenous CSF space^3, 4^. Due to this glymphatic flow, waste solutes travel though meningeal lymphatic system to the outside the brain, which are drained to dCLN^1^. The glymphatic-lymphatic system of CSF influx and ISF efflux show changes with aging and several neurological disorders such as Alzheimer’s disease (AD), subarachnoid hemorrhage, stroke, and traumatic brain injury (TBI)^2, 5^. Glymphatic-lymphatic clearance system maintains homeostasis in the brain, but its disturbance could yet be knowingly manipulated by any method in pre-clinical or clinical studies.

FUS provides ultrasonic energy to a small volume of the target tissues non-invasively. The interaction of FUS and MB that are injected in the blood vessel can stimulate the vascular endothelium to mechanically open the blood-brain barrier (BBB) for several hours^6^ and it has been used as a tool for drug delivery to the brain across the BBB. However, interestingly without any drugs delivery, FUS-MB in AD model reduced the deposition of Aβ and tau, improved memory, and increased internalization of Aβ in microglia^7–9^. These effects were mostly considered to be due to BBB opening, while assuming that brain-resident cells disposed on site within the brain all the abnormal proteins such as A or tau^9, 10^ β. We propose that the FUS-MB shows effects on the waste disposal from the brain parenchyma to CSF and then lymphatics in this investigation. This effect might have improved the behavioral impairment in the previous mouse experiments^7–10^. Based on these therapeutic effects, a single-arm, non-randomized phase 2 trials have been recently performed to evaluate the efficacy of FUS-MB in AD patients^6^. As the applications in clinical practice are important, it is also necessary to investigate the mechanism of the effects.

In this study, we investigated that the cause of Aβ reduction in the whole brain of 5XFAD mice by FUS-MB. FUS-MB decreased the solute Aβ in the brain parenchyma, but not the amyloid plaques, while increasing the amount of Aβ in the CSF. After FUS-MB, clearance of parenchymal Aβ had been enhanced through CSF to dCLN, which was proven to be blocked by lymphatics ligation. The reduction of solute Aβ in the parenchyma by repeated FUS-MB would have led to the inhibition of further plaque formation and consequent maintenance of working memory along aging. FUS-MB would have influenced brain-to-CSF drainage in AD progress in 5XFAD. This simple observation raises the possibility of novel use of FUS-MB for enhancing glymphatic-lymphatic clearance of abnormal protein aggregates in such neurodegenerative diseases as AD.

## Results

### Aβ reduction by FUS-MB in the entire brain further to the target regions

The BBB opening induced by FUS-MB around the ipsilateral hippocampus was confirmed by Evans blue staining (Fig. 1a). FUS-MB in 5XFAD mice reduced amyloid deposition in the contralateral hemisphere as well as in the ipsilateral regions (Fig. 1c). Immunohistochemistry of Aβ on the serial sections of the entire brain of mice treated with FUS-MB (n=6) showed the reduction of total area of Aβ deposits in brain region not directly targeted by FUS-MB (Fig. 1d). As a result of immunostaining with antibodies for microglial marker Iba1 (ionized calcium binding adaptor molecule 1) and reactive astrocytic marker GFAP (glial fibrillary acidic protein), it was shown that FUS-MB reduced gliosis in the hippocampus and entorhinal cortices, contralateral as well as ipsilateral (Fig. 1e). In the wild type mice, microglia activation was also decreased by FUS-MB (Fig. 1e). Therefore, FUS-MB reduced amyloid deposits and ameliorated glial activation in the entire brain.

**Figure 1.**
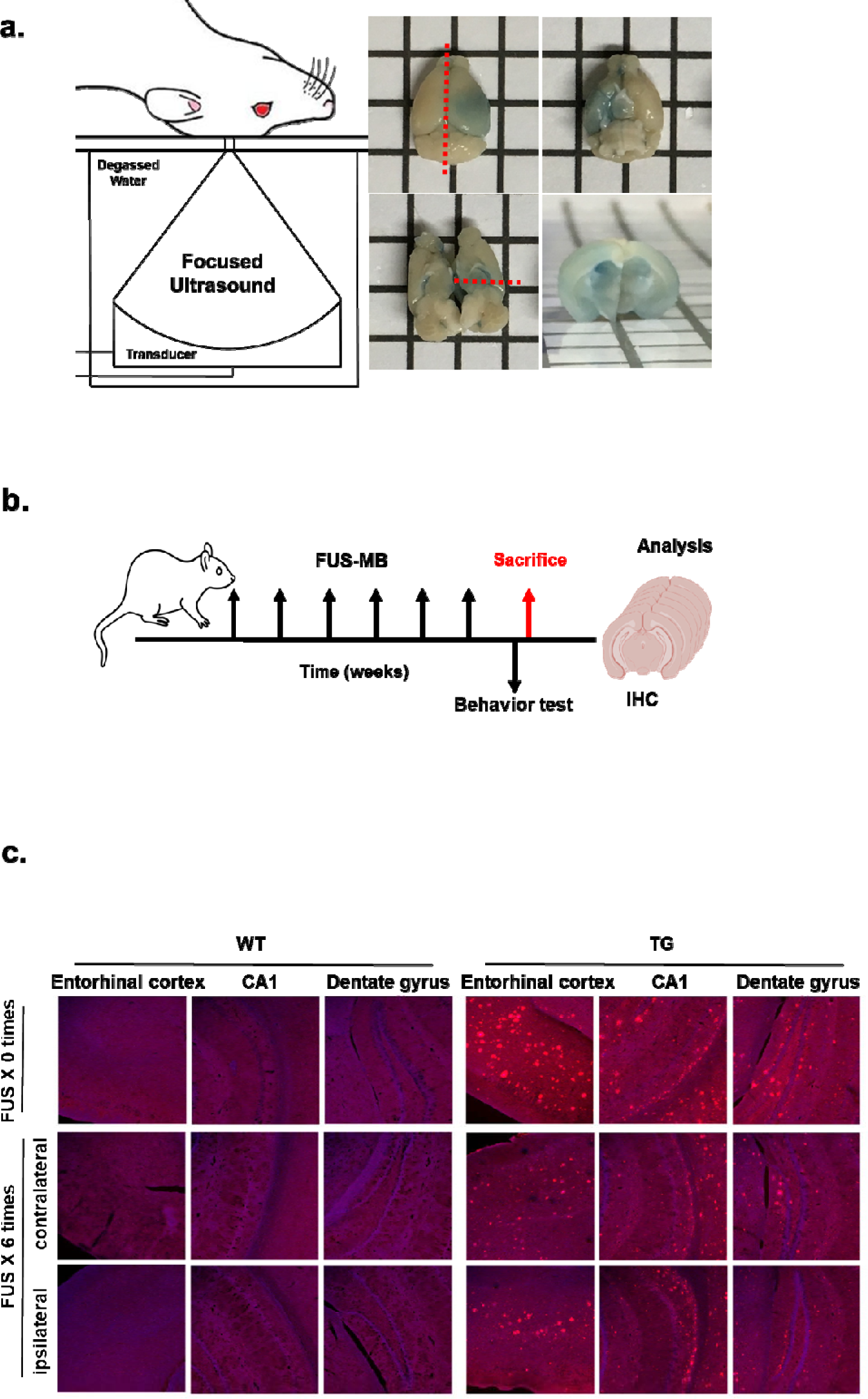
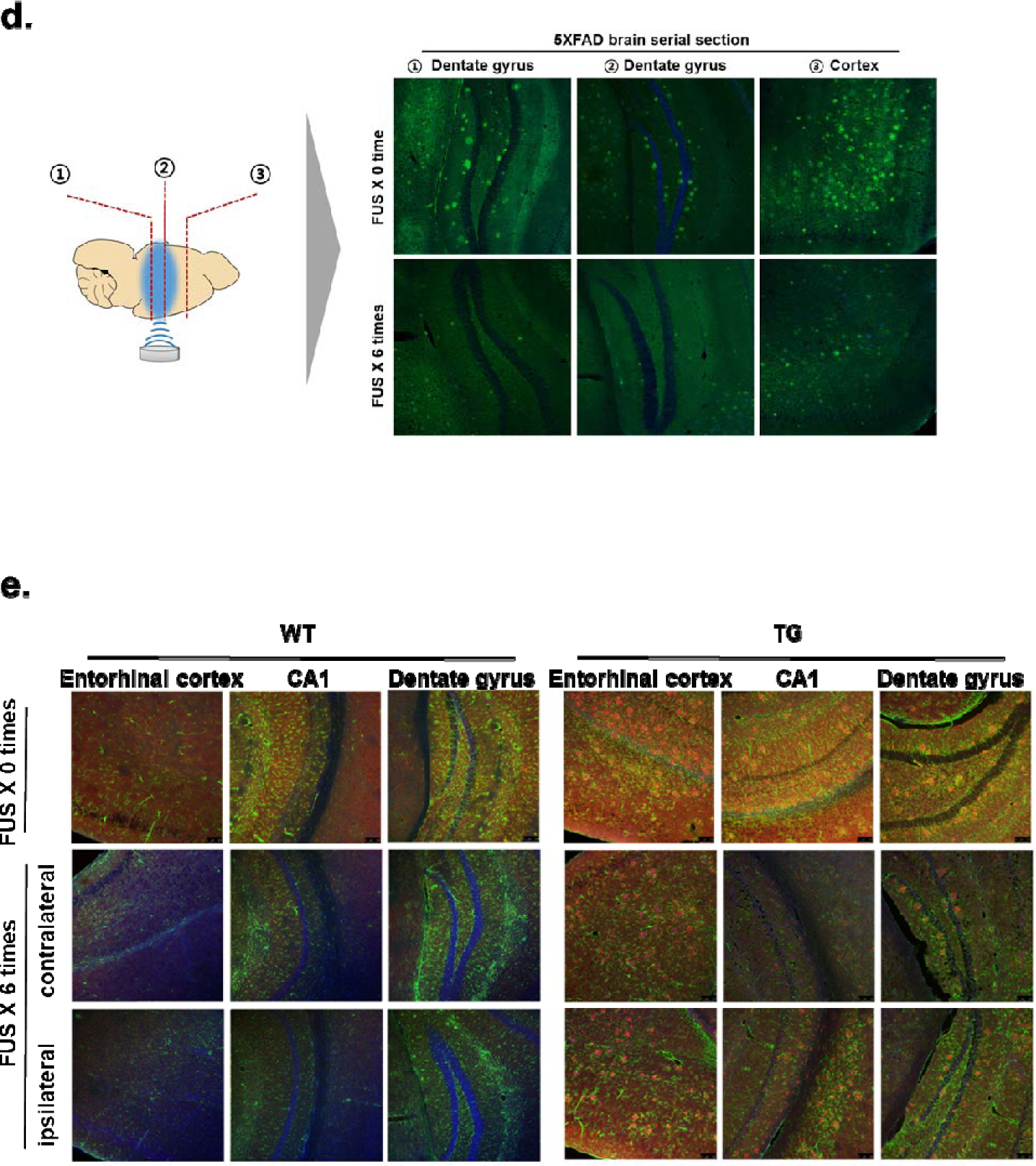
Pathological changes in 5XFAD by repeated FUS-MB. **a.** Schematic image of FUS setup and representative image confirming BBB opening with 2% Evans blue after FUS-MB. The BBB opening region was identified around the ipsilateral hippocampus. The red dashed line represents the cut section. **b**. Design of experiments including 6 repeated FUS-MB, behavioral test and analysis. **c**. Representative images of Aβ immunostaining in the entorhinal cortex and hippocampus of 5XFAD and age-matched control after 6 repeated FUS-MB. **d**. Representative images of Aβ immunostaining in serial sections of 5XFAD brain after FUS-MB. The deposition of Aβ decreased in the entire brain by FUS-MB. **e**. GFAP (green), Iba1 (red), and DAPI (blue) immunostaining images in hippocampus (CA1 and DG) and entorhinal cortex from wild type and 5XFAD mice after 6 repeated FUS to observe changes in astrocyte and microglia activation. Reactive astrocytes were unchanged by FUS-MB, but microglia activation was reduced by FUS-MB.

### FUS-MB improved solute Aβ clearance but not plaques

The lymphatic system in the CNS plays a role in drainage of solute wastes through CSF, and the lymphatic system damage causes amyloid deposition^2^. Ligation of lymphatics to dCLN was performed to reveal its blocking effect upon Aβ and plaques in the entire brain by FUS-MB which increases Aβ clearance to the CSF space. The 6E10 positive Aβ amyloid deposits were significantly decreased by FUS-MB in 5XFAD and dCLN-ligated 5XFAD compared to their controls, respectively (Fig. 2b, c). Thioflavin S (ThS) positive amyloid plaques increased the areas after the lymphatics ligation in dCLN ligation compared to non-ligation group of 5XFAD. ThS positive amyloid plaques did not decrease after FUS-MB in non-liagtion group of 5XFAD (Fig. 2d, e). When the changes after FUS-MB and/or dCLN ligation, FUS-MB decreased Aβ areas but not the areas of amyloid plaques (Fig. 2f). This finding suggested that FUS-MB inhibited plaque formation by decreasing ambient Aβ quantity in the brain and it is probably due to the increased drainage of solute Aβ. These findings support that amyloid plaques are waste depository where the surplus Aβ resided in the brain due to insufficient/ blocked clearance, rather than stopping to go outside of the brain.

**Figure 2.**
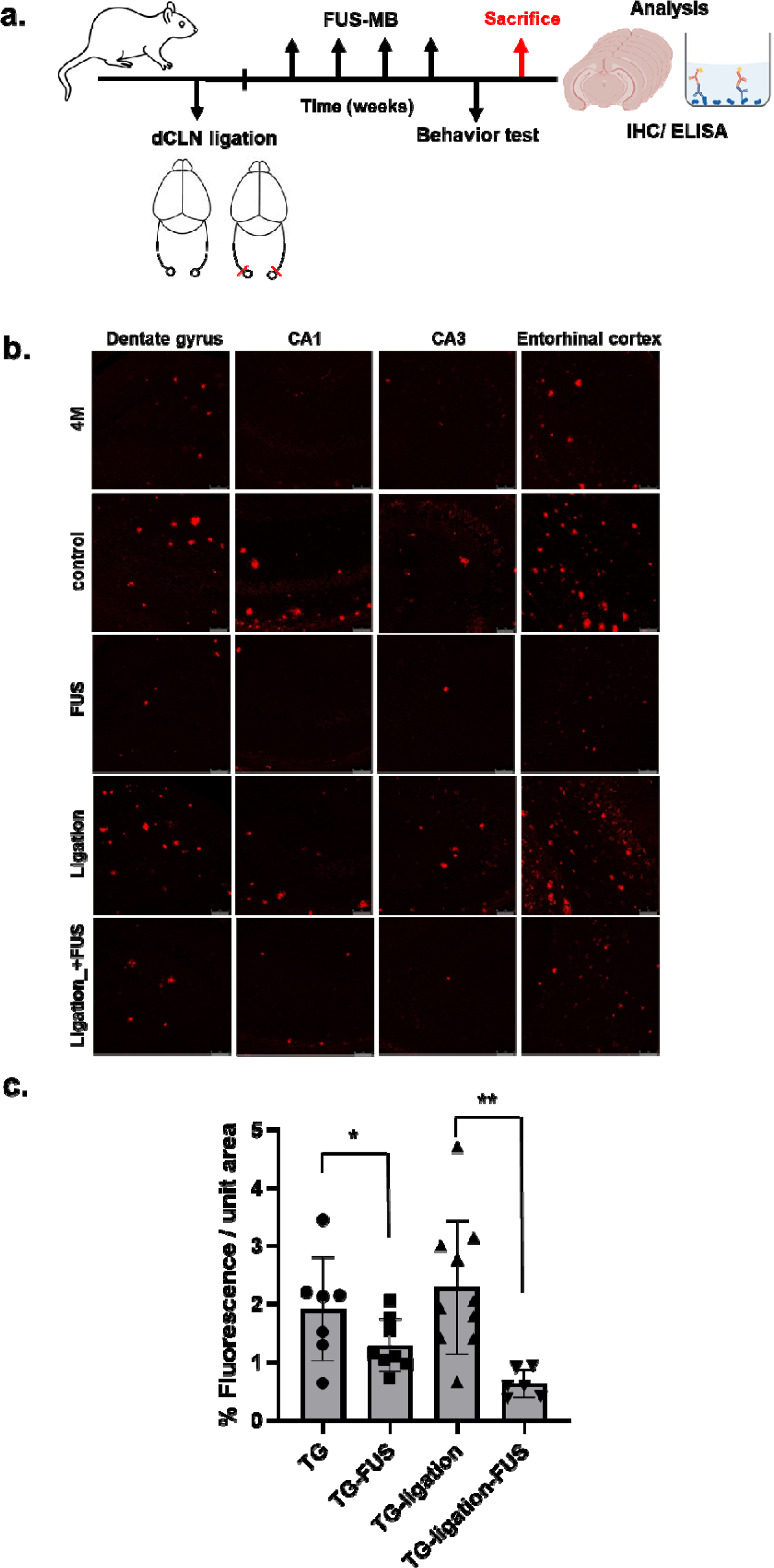
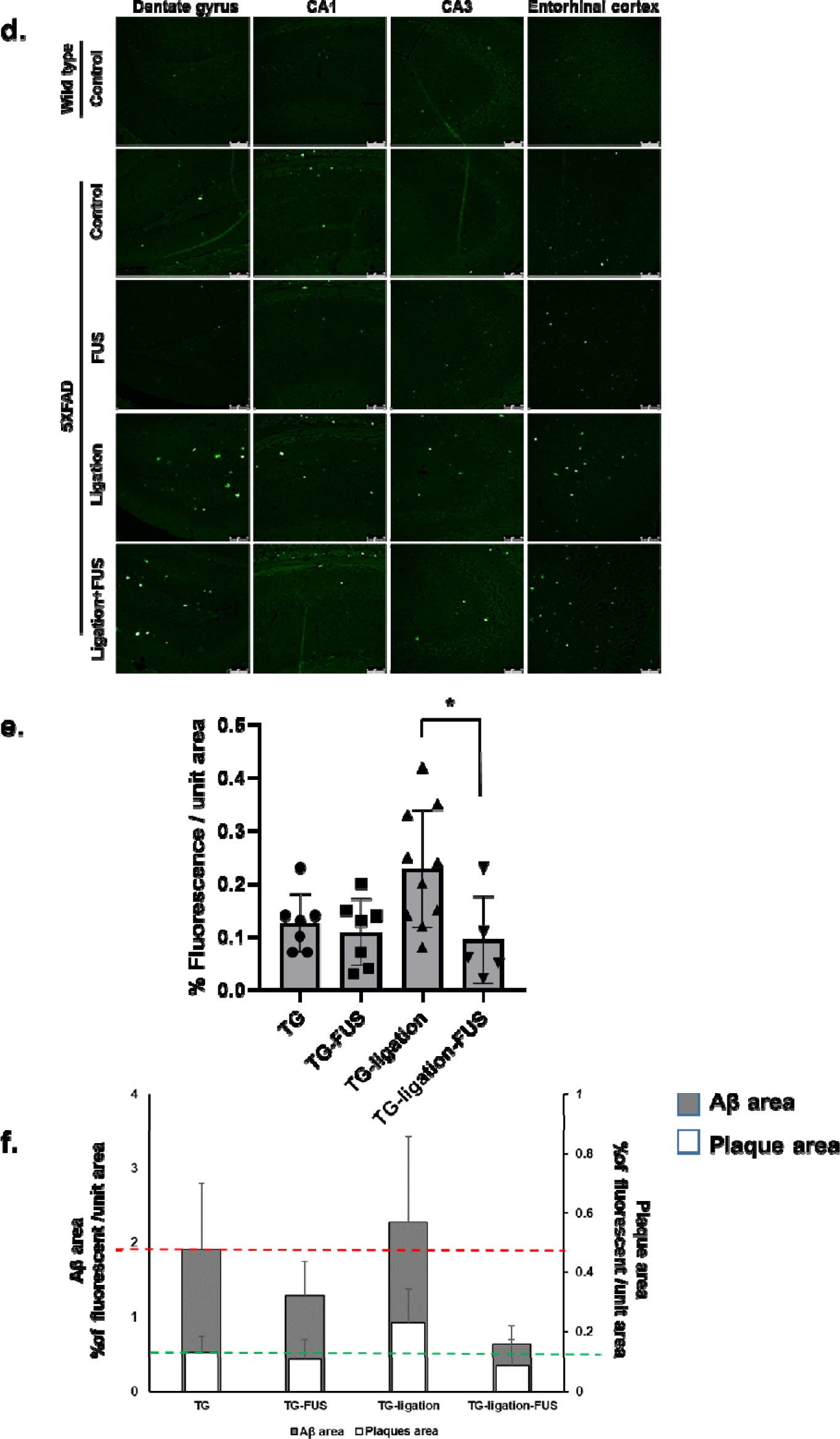
Characterization of Aβ reduction in 5XFAD by repeated FUS-MB. **a** Design of experiments including dCLN ligation, 4 repeated FUS-MB and analysis. **b**. Representative images of Aβ immunostaining in 4 month old mice before the start of the experiment and 6 month old mice after the experiment procedure. **c**. Quatification of the Aβ immunostaining area in the hippocampus and entorhinal cortex. Each dot indicates the quantitative values from 4 sections per mouse (n=6∼10/group). The deposition of Aβ was significantly decreased by FUS-MB. **d**. Representative images of amyloid plaques staining with ThS in 6 months old mice after experiment procedure. **e**. Quatification of the area of plaques in each group. Each dot indicates the average of quantitative values from 4 sections per mouse (n=6∼10/group). The number and area of plaques increased only in the dCLN ligated 5FAD group, and there was no difference in the other groups. **f**. Comparison of mean values of Aβ immunostaining (white) and amyloid plaques (grey). Data were expressed as AVE±SD (*P* < 0.05 (*)).

### CSF Aβ drainage by FUS-MB via meningeal lymphatics

The concentration of CSF Aβ_1-42_ was measured in the 5XFAD mice with or without dCLN ligation to reveal the effect of lymphatic obstruction on CSF drainage of solute Aβ. Compared to 5XFAD without FUS-MB, the concentration of CSF Aβ_1-42_ slightly elevated with FUS-MB in non-ligation group (Fig. 3a). Here, dCLN ligation increased CSF Aβ much further by FUS-MB (Fig. 3a). This indicates that CSF Aβ_1-42_ are cleared through meningeal lymphatics draining to dCLNs. and On immunohistochemistry, while dCLN ligation showed rare Aβ around meningeal vessel, FUS-MB induced the prominent accumulation around the meningeal arterial and venous. This accumulation was exaggerated by FUS-MB in dCLN ligation group (Fig. 3b). It was interpreted that solute Aβ were snapshot while stuck in around vascular spaces due to the back-pressure of lymphatic outflow obstruction. However, it is not certain that Aβ are just leaving or staying in the neighbor of meningeal vessels.

**Figure 3.**
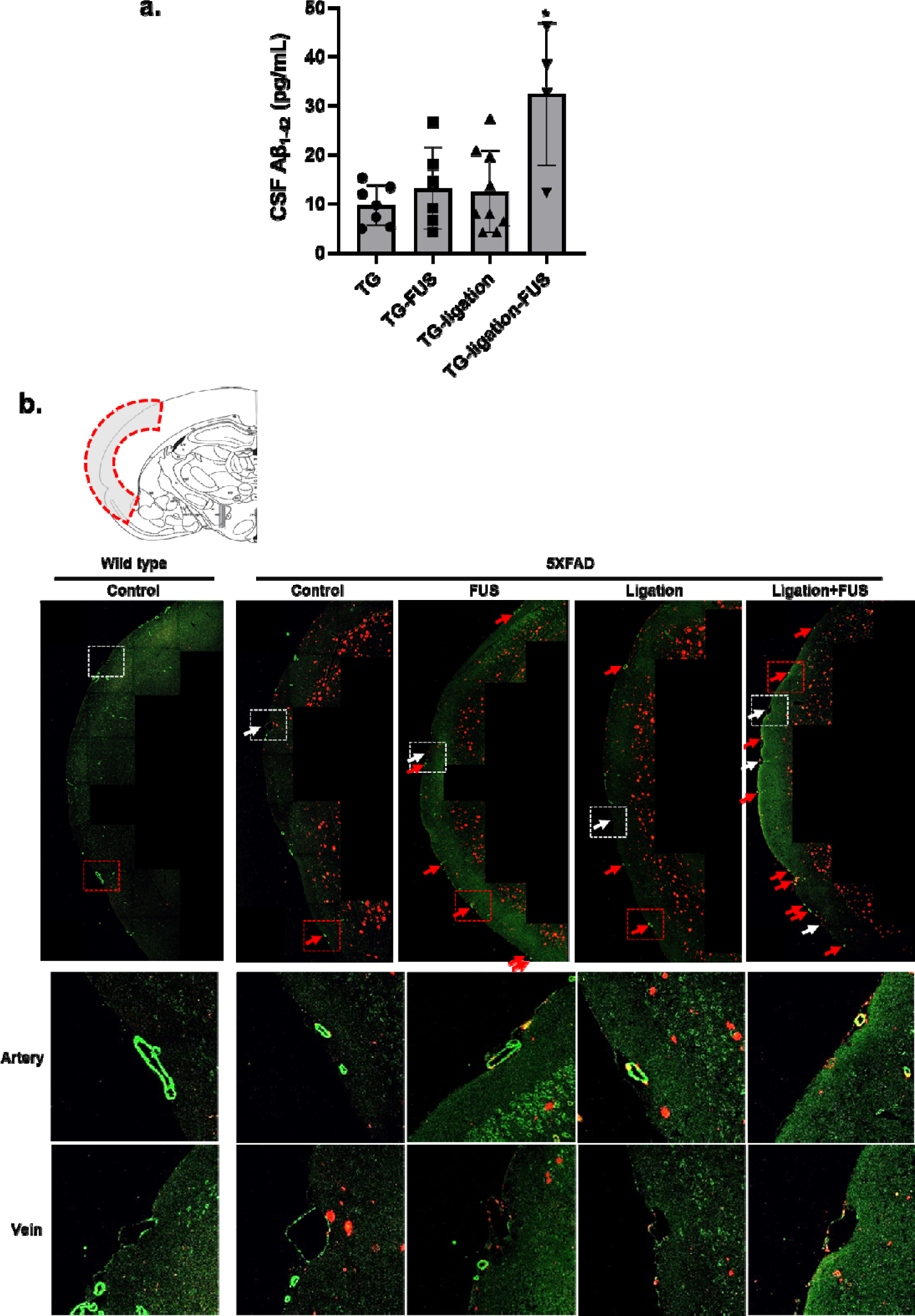
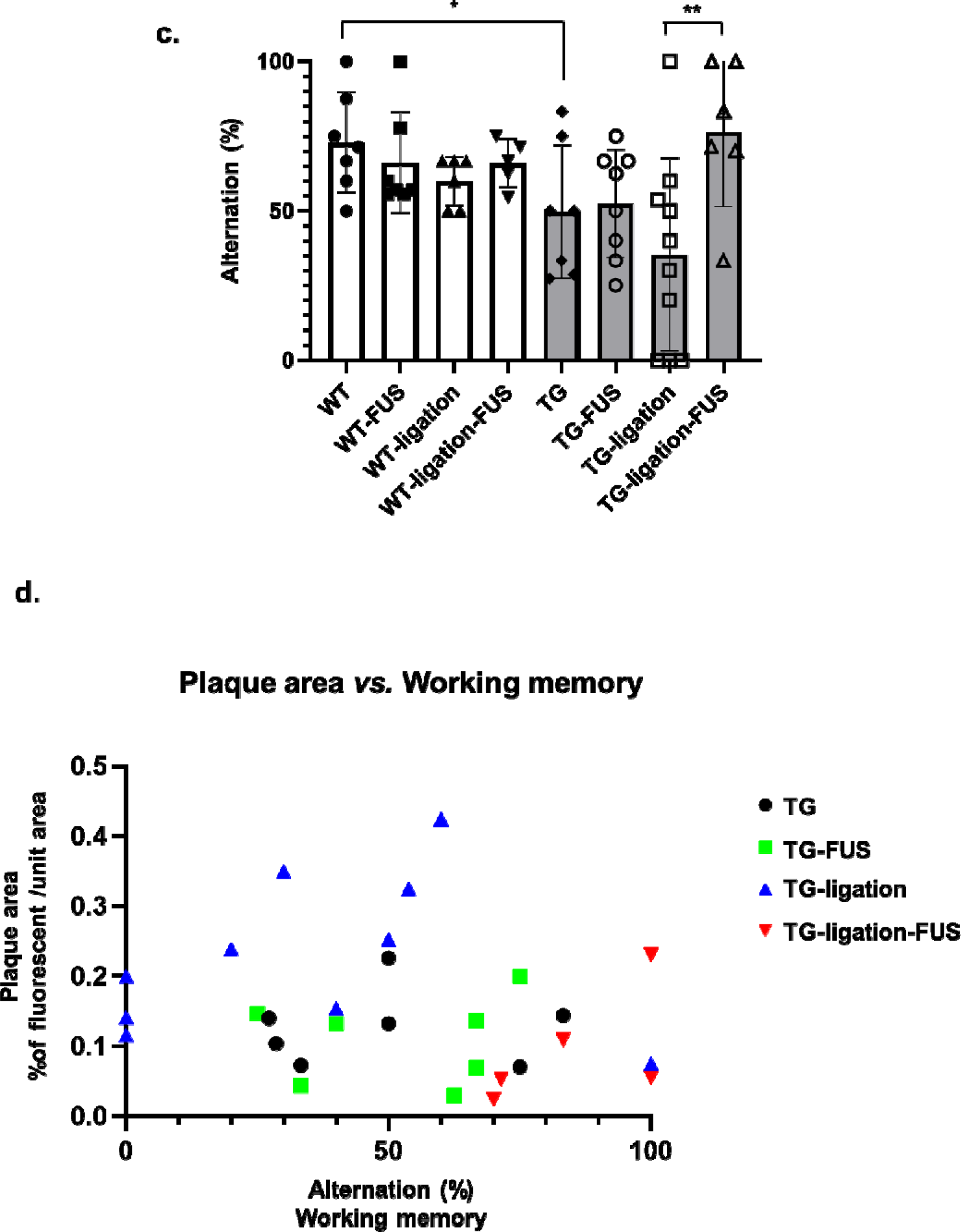
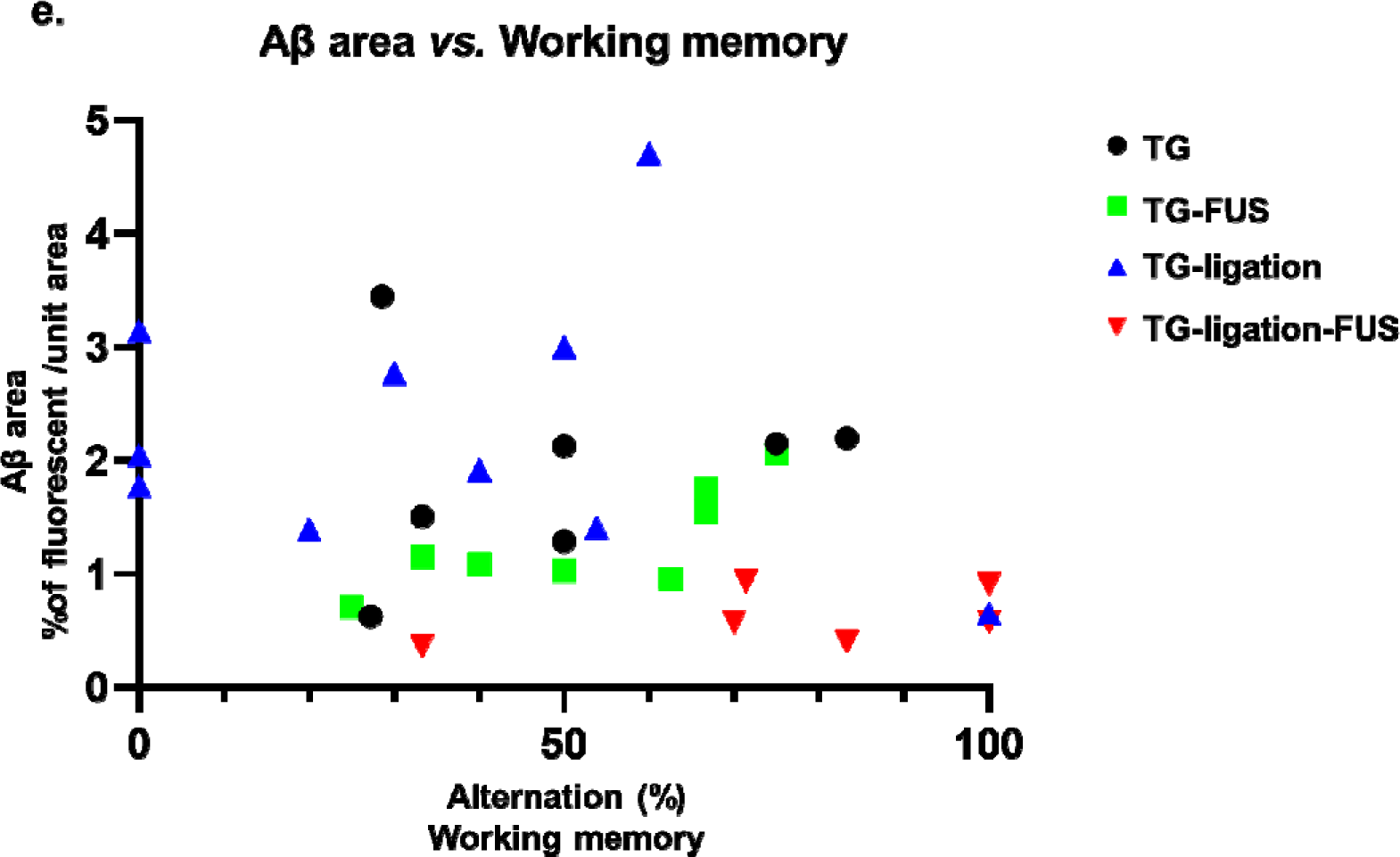
Cognitive changes and CSF drainage of Aβ by repeated FUS-MB The concentration of CSF Aβ. **a.** was measured in 5XFAD groups with ELISA. The concentration of CSF Aβ significantly increased in dCLN ligated 5XFAD after FUS-MB. Representative images of Aβ (red) and smooth muscle actin (SMA, green) immunostaining in the meningeal vascular region. Red arrows indicate the presence of amyloid around artery. And white arrows indicate the presence of amyloid around vein. The red and white dashed boxes are the artery and vein regions, respectively. They are magnified beow. An increase of amyloid around meningeal vessels was observed especially in dCLN ligated 5XFAD after FUS-MB. **c**. Measurement of changes in spatial working memory with Y-maze test. After FUS-MB treatment, spatial working memory of 5XFAD tended to be improved and it was observed that cognitive function was not impaired in the dCLN ligated 5XFAD after FUS-MB. **d**. Correlation between spatial working memory and area of Aβ immunostaining in all 5XFAD groups (n= 13). **e**. Correlation between concentration of CSF β1-42 and the area of Aβ immunostaining in dCLN ligated 5XFAD groups (n= 6). Area of amyloid immunostaining was correlated with spatial working memory or CSF Aβ. Data were expressed as AVE±SD (*P* < 0.05 (*), *P*<0.005 (**)). β1-42

### Working memory by dCLN ligation and/or FUS-MB in 5XFAD mice

By dCLN ligation, 5XFAD mice suffered from the impairment of working memory, which was significantly improved by FUS-MB (Fig. 3c). Plaque burden, represented by ThS stained area (ThS area), was not correlated (*p*=0.37,n=29) with the alternation % of the Y-Maze tests (Fig. 3d, Supplementary Fig. 2). However, there was the tendency that ThS area was lower in dCLN ligation with FUS-MB than in dCLN ligation only group. The ThS area was positively correlated (*p*=0.0005, n=29) with 6E10 positive Aβ amyloid deposits (Aβ area), and unlike the ThS area, Aβ area tended to be negatively correlated (*p*=0.15, n=31) with alternation % of Y-Maze (Fig. 3e, Supplementary Fig. 2). CSF Aβ was not correlated with ThS area, Aβ area, or Y-Maze scores, higher the CSF Aβ better the alternation % of Y-Maze, especially in the FUS-MB treated dCLN ligation model (Supplementary Fig. 2). Taken together, FUS-MB prevents cognitive impairment by enhancing the drainage of solute Aβ drainage through CSF-lymphatics system.

### Changes of brain cells by FUS-MB and/or dCLN ligation

The effects of FUS-MB on brain cells were examined to evaluate the changes on the cells which could be caused directly by FUS-MB or indirectly by the pathological changes. After dCLN ligation in entorhinal cortex, H&E and TUNEL staining revealed the loss of neurons which was rescued by FUS-MB (Fig. 4a). Also, GFAP positive astrocytes increased and FUS-MB prevented this increase. In other areas such as dentate gyrus, CA1 and CA3 in hippocampus, no increase after dCLN ligation and no rescue by FUS-MB was observed (Fig. 4b). Propensity of Iba1 positive microglia increased after dCLN ligation in entorhinal cortex, which was also prevented by FUS-MB (Fig. 4c). However, there was no prominent difference in other areas. Microglia showed up surrounding the plaques and their decrease was accompanied by the decrease of 6E10 positive Aβ deposits by FUS-MB (Fig. 4c). FUS-MB was not found to induce neuronal loss or reactive astrocytes and microglial propensity surrounding Aβ deposits, but to prevent neuronal loss and activation of glial cells induced by dCLN ligation in 5XFAD AD mouse model.

**Figure 4.**
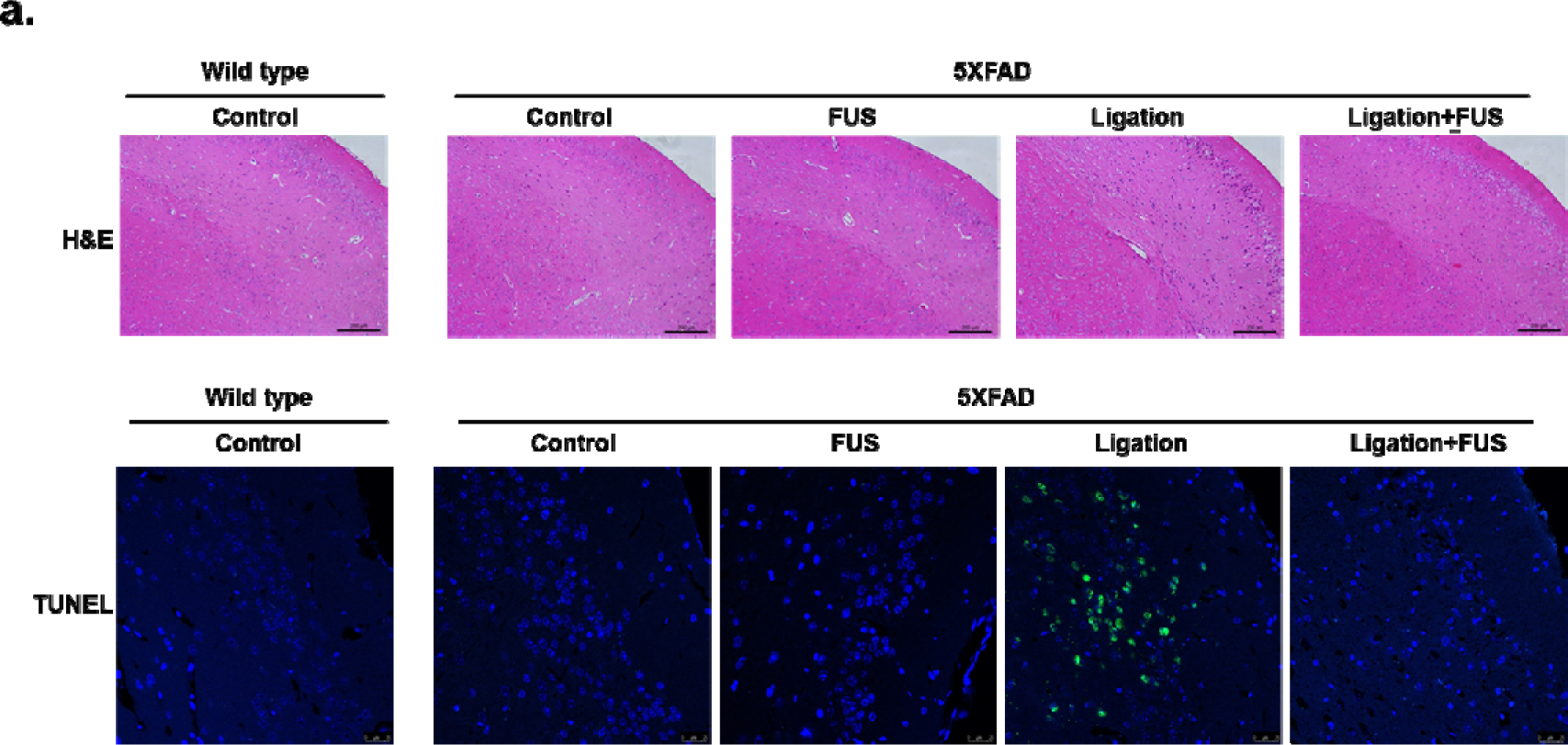
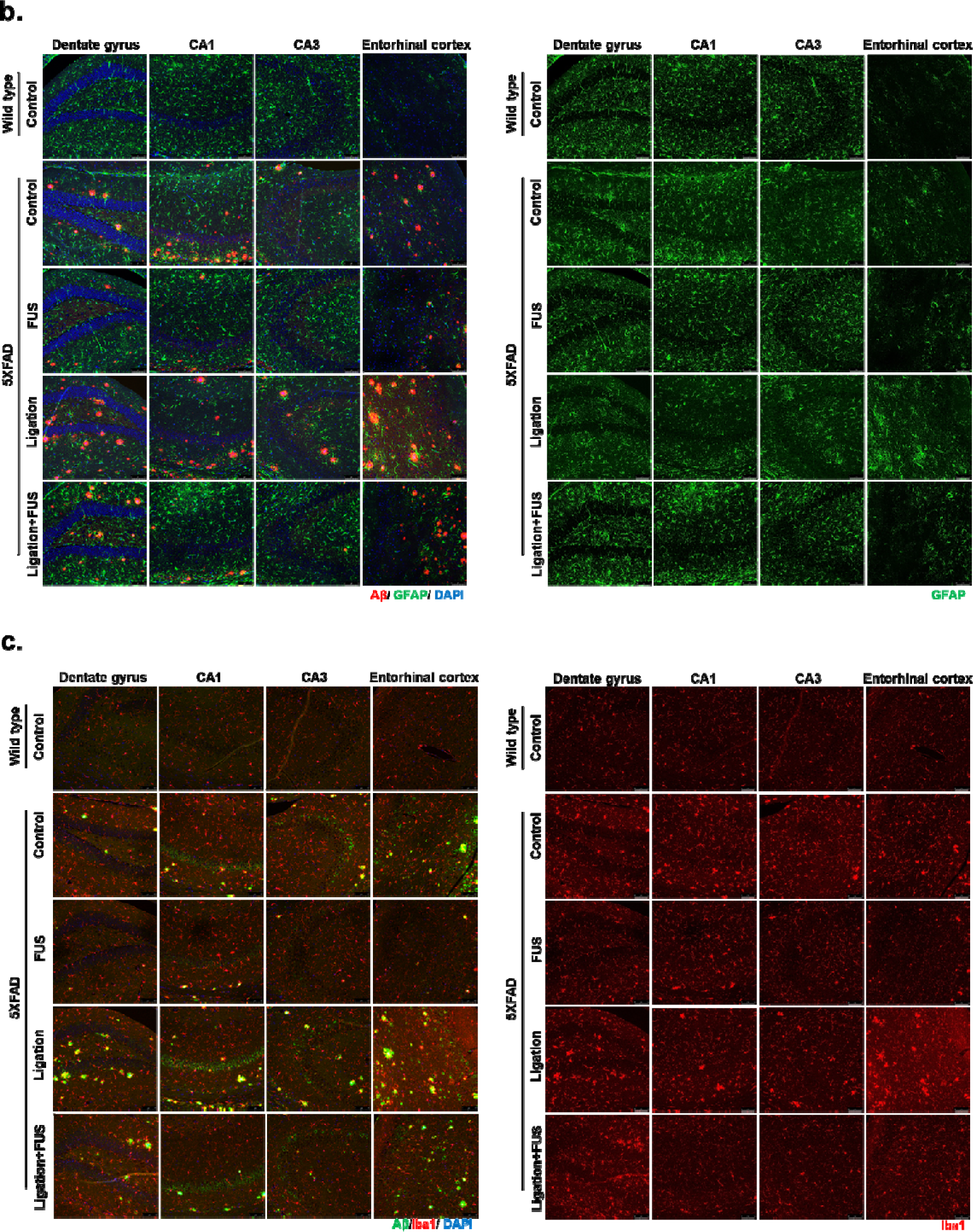
Effects of brain cells in FUS-MB treated 5XFAD. **a** Representative images of neuronal loss in the entorhinal cortex by H&E and TUNEL staining. Neuronal damage was observed in the entorhinal cortex of dCLN ligated 5XFAD, but no neuronal damage was observed in dCLN ligated 5XFAD after FUS-MB. **b**. Representative merged images of GFAP (green), Aβ (red), and DAPI (blue) immunostaining in hippocampus and entorhinal cortex (left). The isolated images of GFAP to compare the changes of reactive astrocytes (right). Reactive astrocytes were reduced only in entorhinal cortex after FUS-MB. **c**. Representative merged images of Iba1(red), Aβ (green), and DAPI (blue) immunostaining (left). The isolated images of Iba1 to compare the changes of microglia activation (right). Microglial activation was reduced in the entire brain by FUS-MB.

**Figure 5.**
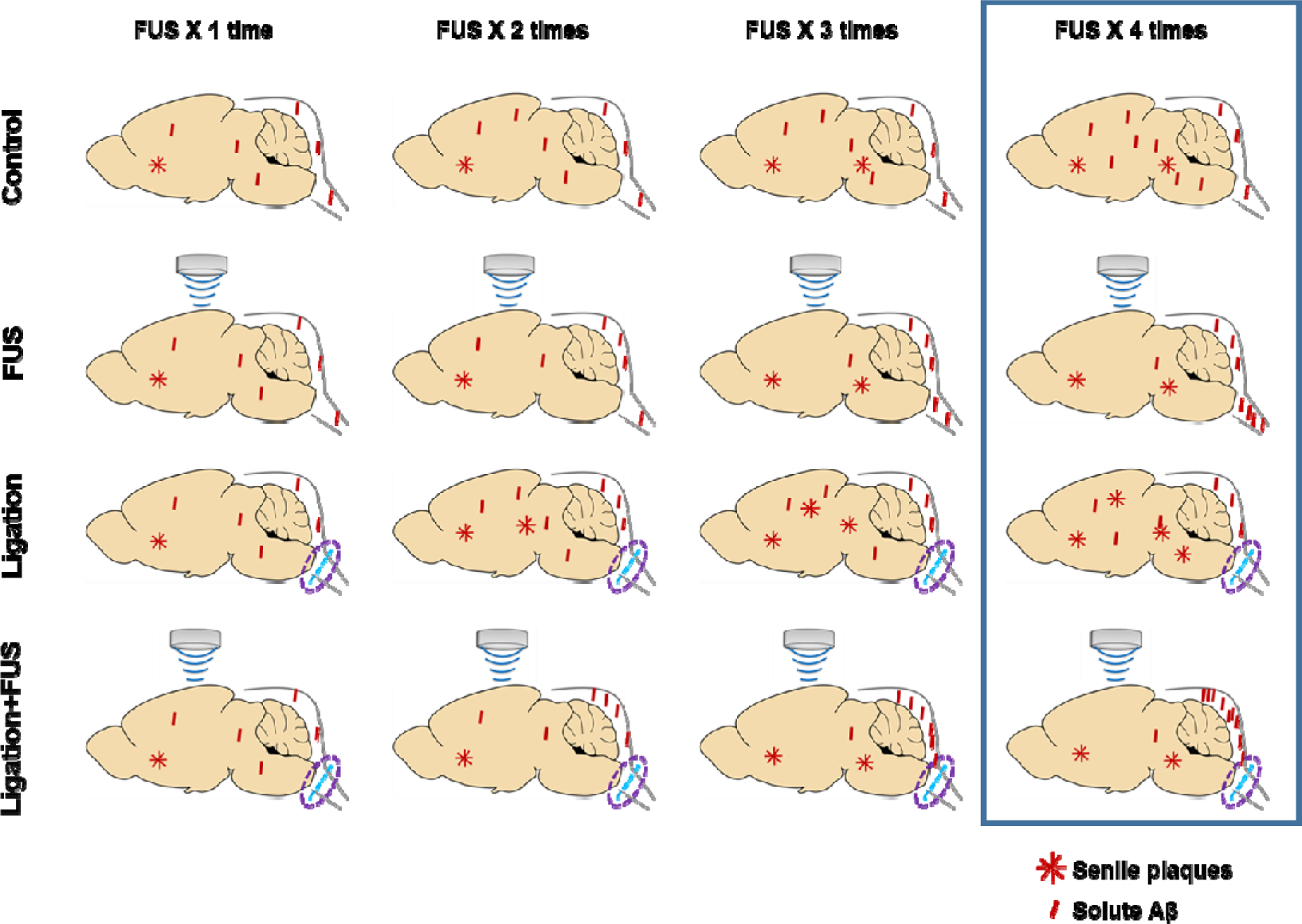
Preventive effect of FUS-MB through CSF drainage in AD model Schematic image of solute Aβ clearance and plaque deposition during repeated FUS (1∼ 4 times). Aβ and plaque deposition increased in 5XFAD mouse for 2 months. The dCLN ligation group showed increased plaque formation without increasing solute Aβ deposition compared to 5XFAD group. FUS-MB treated groups showed the reduction of solute Aβ in the parenchyma and no changes of plaques area. Reduction of soluble Aβ shifted to CSF and maintained cognitive function.

## Discussion

In this study, FUS-MB improves Aβ clearance and cognitive function in AD model by glymphatic-lymphatic drainage. This decrease of Aβ deposits was observed in the contralateral and ipsilateral regions remote from the focused regions by FUS-MB in a series of sections (Fig. 1c, d). The overall reduction of Aβ deposits by FUS-MB raises the question whether FUS-MB decrease Aβ production or stimulate Aβ clearance. While FUS-MB decreased overall Aβ deposits at 4 months before FUS-MB (Fig. 2b), the Aβ concentration in CSF tended to increase after ligation and increased much more after FUS-MB in 5XFAD mice after dCLN ligation (Fig. 3a). We interpret this indicates that FUS-MB effectively removed Aβ effectively upon our protocol (Fig. 1c, d). Differential Aβ clearance from amyloid plaques or Aβ deposits was examined using ThS staining and IHC with 6E10 antibody recognizing N-terminal region of Aβ 40 and Aβ 42 isoforms and their precursor forms (Fig. 2). In naïve 5XFAD and dCLN ligation groups, Aβ deposits were similar and FUS-MB decreased much of the Aβ burden (Fig. 2b, c). In contrast, ThS stained plaques were not different between naïve and FUS-MB treated 5XFAD mice (Fig. 2d, e). After dCLN ligation, plaque burden variably increased and FUS-MB decreased the burden in their area (Fig. 2d, e). These results indicated that FUS-MB cleared solute Aβ but not the plaques (Fig. 2f), while ligation blocking lymphatic drainage resulted in the increase of plaques area (Fig. 2d, e). Multivariate regression to explain behavior on Y maze tests did not disclose any explanatory variable, while ThS areas and Aβ areas were correlated significantly with r square of 0.38 and no other significant correlation between CSF Aβ concentration, ThS areas and Aβ area (Supplementary Fig. 2a). The only finding to explain the variability of memory performance on Y maze was the fact that it was working memory after FUS-MB in dCLN ligated groups (Fig. 3c). Despite the amount of sustaining plaques similar to those of naïve 5XFAD, after FUS-MB treatment in dCLN ligation compared with naïve 5XFAD, the amount of plaques was significantly smaller than dCLN ligation (Fig. 3d, e). Taken together, FUS-MB could enhance solute Aβ drainage from brain parenchyma, and improvement of working memory in FUS-MB, exaggerated in lymphatic ligation model, was correlated with solute Aβ clearance and plaques formation in the brain.

The interaction of FUS with MB may induce various outcomes in the targeted brain, ranging from the originally proposed BBB opening to the induction/facilitation of Aβ clearance. Probable mechanisms of increased disposal of solute wastes by FUS-MB are 1) Enhanced clearance of waste by degradation on site. Microglia was activated and took up Aβ and tau by phagocytosis after FUS-MB and it also induced immune cells.^7, 9^ MRI guided FUS reduced plaque load at the targeted brain cortex, while IgG and IgM decorated around the plaques and the role of endogenous antibodies was suggested.^11^ 2) Enhances clearance through BBB through transporters like RAGE or LRP1 which regulates transport of Aβ. The expression of RAGE is upregulated and that of LRP1 is downregulated on the BBB in AD patients^12^.

RAGE1 and LRP1 regulated Aβ levels in the brain and plasma^13^. Vascular smooth muscle actin was lost and smooth muscle cells degenerated in AD and these changes might be associated with decreased clearance through BBB.^14^ FUS are yet to be found to influence these molecules. 3) Enhanced clearance of Aβ by glymphatic-lymphatic system. Aβ clearance increased in relation with non-REM sleep. Impairment of meningeal lymphatics induced amyloid accumulation, and AAV1-CMV-mVEGF-C treatment enhanced CSF drainage and meningeal lymphatics function and improved spatial learning and memory.^2^ As water channel aquaporin-4 (AQP4) in astrocytic end feet plays a role in glymphatic flow and maintained CSF influx in *Aqp*4 knockout mice^15^. In AD, AQP4 were expressed abnormally and mislocalized.^16^ FUS can also affect AQ4 and facilitate waste disposal by stimulating glymphatic-lymphatic system.

We decided to test the contribution of glymphatic-lymphatic system to explain the effect of FUS-MB on Aβ clearance. CSF Aβ concentration tended to increase in some animals after dCLN ligation, however, FUS-MB increased much CSF Aβ after dCLN ligation (Fig. 3a). FUS-MB was sure to dispose solute Aβ through the vascular spaces into meningeal lymphatic vessels (Fig. 3b). In other words, dCLN ligation disclosed the effect of FUS-MB upon glymphatic-lymphatic clearance. Interestingly, dCLN ligation resulted in the aggravation of memory impairment, but individual variation obscured the significant difference (Fig. 3c). The beneficial effect of FUS-MB on memory dysfunction in naïve 5XFAD was also seen but with much variation. However, the effect of FUS-MB on dCLN ligation was prominent and thus significant (Fig. 3c). dCLN ligation and consequent meningeal lymphatic blockage exaggerated the effect of FUS-MB on glymphatic-lymphatic system (Fig. 3c). We suspect that the variation of memory impairment in 5XFAD syngeneic mice along their aging (4 months old to 6 months old) might be related, at least partially, with the normalcy of lymphatic clearance capacity in individual mice. Taking advantage of dCLN ligation model and measurement of CSF Aβ and also the memory scoring on Y maze test, we now demonstrate that FUS-MB increased glymphatic clearance of Aβ to CSF spaces through meningeal arteries and/or veins and meningeal lymphatic clearance of Aβ to dCLN (Fig. 3a-c). This effect of FUS-MB on memory function was evident in lymphatic ligation cases with the highest CSF Aβ concentrations (Fig. 3a, c). Which cells or molecules are directly or indirectly influenced by FUS-MB are to be unraveled.

Glymphatic-lymphatic clearance consists of CSF-ISF interface for influx and efflux of toxic solutes such as Aβ and CSF-lymphatic drainage of Aβ. In order to enhance CSF-ISF exchanges, several physiological factors were already presented^17^: 1) increased CSF production, 2) modulated pulsatility of arterial vessels, 3) increased expression of AQP4 water channels in astrocytic end feet, and 4) temporal expansion of the space between cells by acute shrinkage of brain cells. And this last mechanism was related with non-REM sleep during which period, glial cells shrank their sizes down to 60%^18^. With our results, we can say that the effect of the interaction between FUS and MB might be the main contributor to the enhanced clearance on Aβ to cross ISF-CSF border. After intravenous administration of MB, MB circulates though the heart and lungs into the brain arteries, and MB cavitation in the arteries might function to mimic and enhance arterial pulsatility on driving the ISF-CSF efflux of Aβ solutes. The importance of arterial pulsatility modulating the dynamics of CSF-ISF influx was reported using partial artery occlusion, adrenergic agonist dobutamine, and anesthesia ^3, 19^.

Our results demonstrate that glymphatic-lymphatic system is involved and considering differential effect of FUS-MB on plaques and solute Aβ, it might be preferred that FUS-MB acts on solutes Aβ but not plaques (Fig. 2). As the quantity of Aβ and plaques are expressed by Image J-derived areas and thus correlated with each other (38% of mutual explanatory power), the changes of plaques might (or might not) be observed in similar or dissimilar experiments of FUS. We propose that solute Aβ is the target of removal by glymphatic-lymphatic system, deterred by dCLN ligation and facilitated by FUS-MB. Thus far, we do not understand exactly 1) whether CSF is drained mainly through the basal meningeal lymphatic vessels or evenly through dorsal/basal and spinal meningeal lymphatic vessels and 2) how much fraction of CSF is effluxed to each type of meningeal lymphatic vessels. According to the recent report, basal meningeal lymphatic vessels were proposed to the main routes of disposal of CSF to lymphatic vessels, however, they used cisternal injection of MRI contrast agents or even quantum dots. The preferred routes of choosing basal meningeal lymphatic vessels might be taken granted by the basal cistern, the injection site. Therefore, various experimental results, such as injection site and dosage, will enlighten the contribution of the location/type of meningeal lymphatic vessels. Understanding this CSF-lymphatic clearance is still primitive and more facts are expected to be elucidated.

We administered FUS with MB targeted not to a small region, but to almost one-third of a hemisphere (Fig. 1a), and we examined the effects of FUS-MB upon the brain-residing cells regarding gliosis and neuronal loss (Fig. 4). After FUS, acute inflammation is known to take place with glial cell activation decreasing within 24 hours^20^. In this experiment, we examined gliosis a week after 4- or 6-times repeated FUS-MB and observed the decrease of overall decrease of Iba1 positive microglia in entire brain including dentate gyrus, CA1/CA3 hippocampus and entorhinal cortex compared to naïve 5XFAD (Fig. 4c). FUS-MB also decreased GFAP positive activated astrocytes uniquely in entorhinal cortex after FUS-MB and did not in the other brain regions (Fig. 4b). It is not yet known whether the decrease of gliosis regionally by activated astrocytes or globally by microglia after FUS-MB was due to its effect on microglia to decrease phagocytosis or due to the effect on Aβ reduction causing decrease of microglia and activated astrocytes. The effect of FUS-MB on neuronal death only appeared after dCLN ligation. Despite dCLN ligation, FUS-MB might have relieved harmful effect of Aβ on neurons, and naïve and FUS-MB treated 5XFAD mice did not reveal neurons dead (Fig. 4a). We applied FUS-MB for wider areas than the previous FUS-MB BBB opening studies. Nevertheless, the neuronal death is expected to be caused by solute Aβ which was successfully drained by FUS-MB even after dCLN ligation.

In mice we used a model to facilitate efflux of harmful toxic solute which was supposed to be Aβ by FUS-MB and a model to block the drainage through meningeal lymphatics by dCLN ligation. The use of these models improved the contrast of the findings, and we could delineate the importance of improved amyloid clearance and restored memory in AD model mice by enhancing glymphatic-lymphatic clearance. In humans, phase 1 study showed no group differences on amyloid plaque burden on amyloid PET images after repeated FUS in five patients with AD ^21^. Though, this negative result can be simply attributed to the difference between mouse and human^22^, there might be the confounders like the status of lymphatic drainage and many more factors affecting the progress of AD in humans. Roles of added MB were also suggested for effective FUS as MB were distributed in perivascular space, subarachnoid space, and surrounding large veins in patients with AD and ALS and thus might be assumed to have regulated glymphatic efflux by FUS^23^. We also consider the possibility that solute Aβ reduction through Aβ clearance would contribute only to the prevention during preclinical stage rather than established AD or the need for long-term follow-up to observe the preventive effect. Aβ burden should have been removed by FUS-MB before they had injured permanently the neurons and other brain-resident cells.

Taken together, glymphatic-lymphatic system of the brain plays a direct role in Aβ amyloid deposition and drainage, while FUS-MB restored memory by increasing amyloid drainage. The structural changes of the meningeal lymphatic vessels with aging result in the functional consequences of the glymphatic-lymphatic clearance of Aβ and other possible toxic solutes^24^, which is why ageing has become a major risk factor for AD. FUS-MB is a potential preventive measure against dysfunction of glymphatic-lymphatic system in prodromal or early AD. Furthermore, the effect of the current amyloid removal therapeutics, such as antibody therapy, might be enhanced by FUS-MB and should be accompanied or accompanied by clearance of toxic solutes already present or newly generated in the aging brain by antibody therapy. We emphasize that physiologic disposal from the brain parenchyma of the toxic solute macromolecules has the novel preventive and therapeutic possibility for neurodegenerative diseases associated with abnormal proteins such as tau, α-synuclein and TDP43 as well as Aβ.

## Acknowledgments

We appreciate the critical reading by Dr. Min Seok Suh. And this research was supported by the National Research Foundation of Korea (NRF) grant funded by the Korean Government (MSIP) (No.2015M3C7A1028926 and No. 2017M3C7A1048079), and NRF grant funded by the Korean Government (No. 2017R1D1A1B03032037).

## Author contributions

Y.C., E.J.P. D.S.L. designed the study. Y.E., Y.C., S.K., K,H. performed experiments and data analyses. E.J.P. and J.Y.L. contributed experiments for focused ultrasound. Y.C., Y.E. and D.S.L. wrote the paper. All authors read and approved the final manuscript.

## Extended data

**Supplementary Figure 1.**
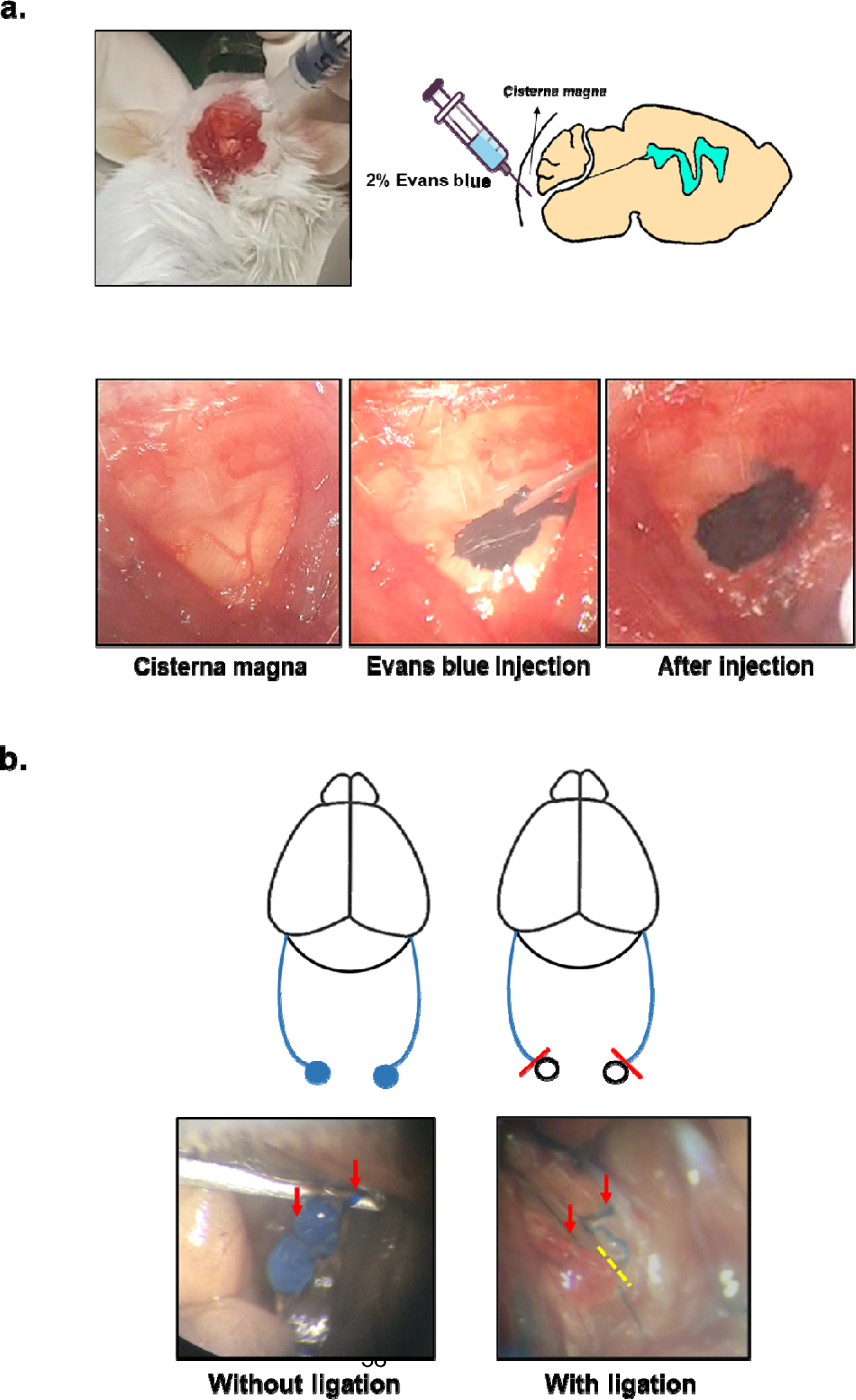
Validation of dCLN ligated animal model **a.** Evans blue was injected into cisterna magna of mice to confirm the ligation of lymphatics. **b.** 30 minutes after Evans blue injection into cisterna magna, CSF and dCLN changed to blue. On the other hand, after lCLN ligation, dCLN did not turn blue.

**Supplementary Figure 2.**
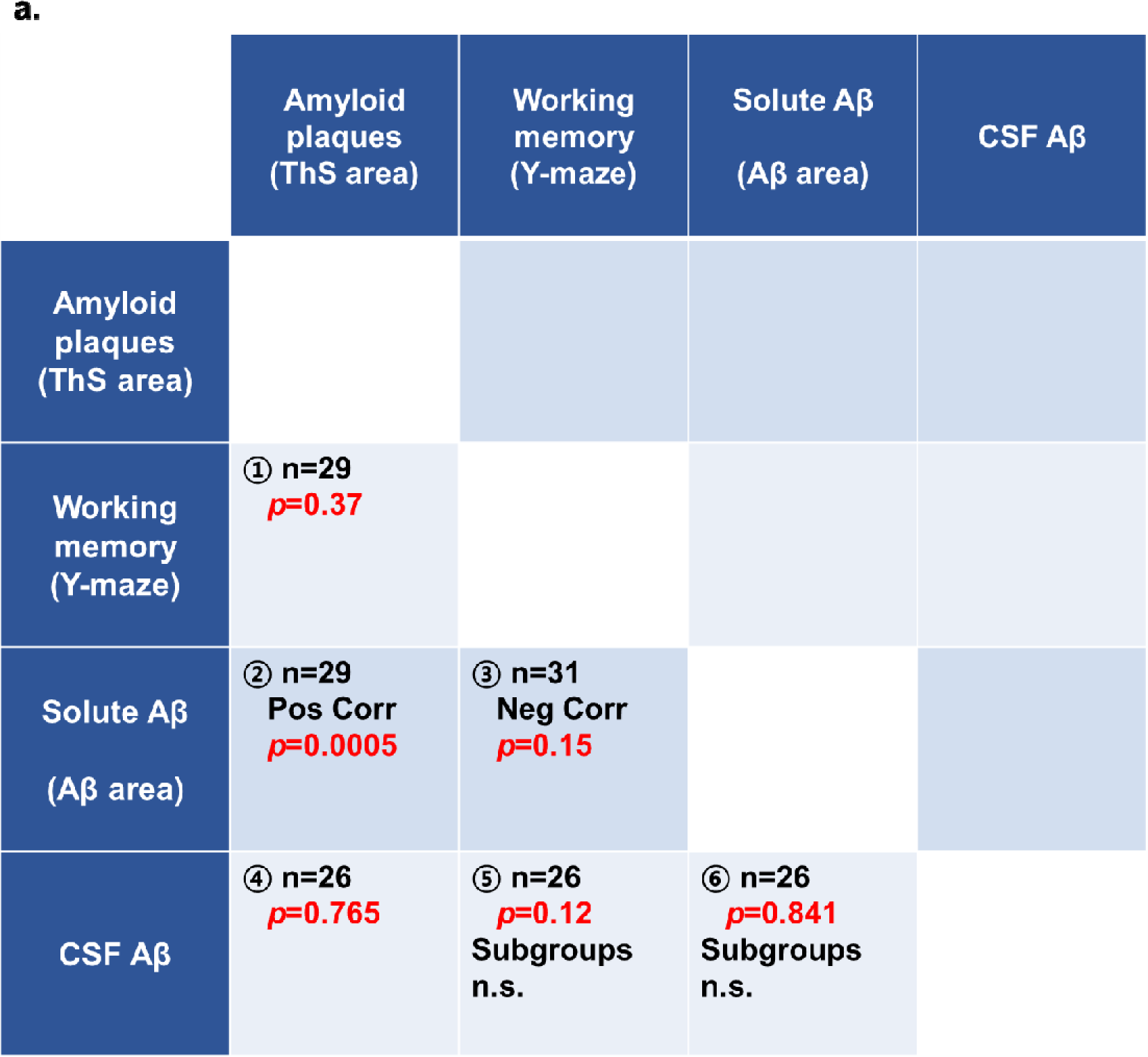
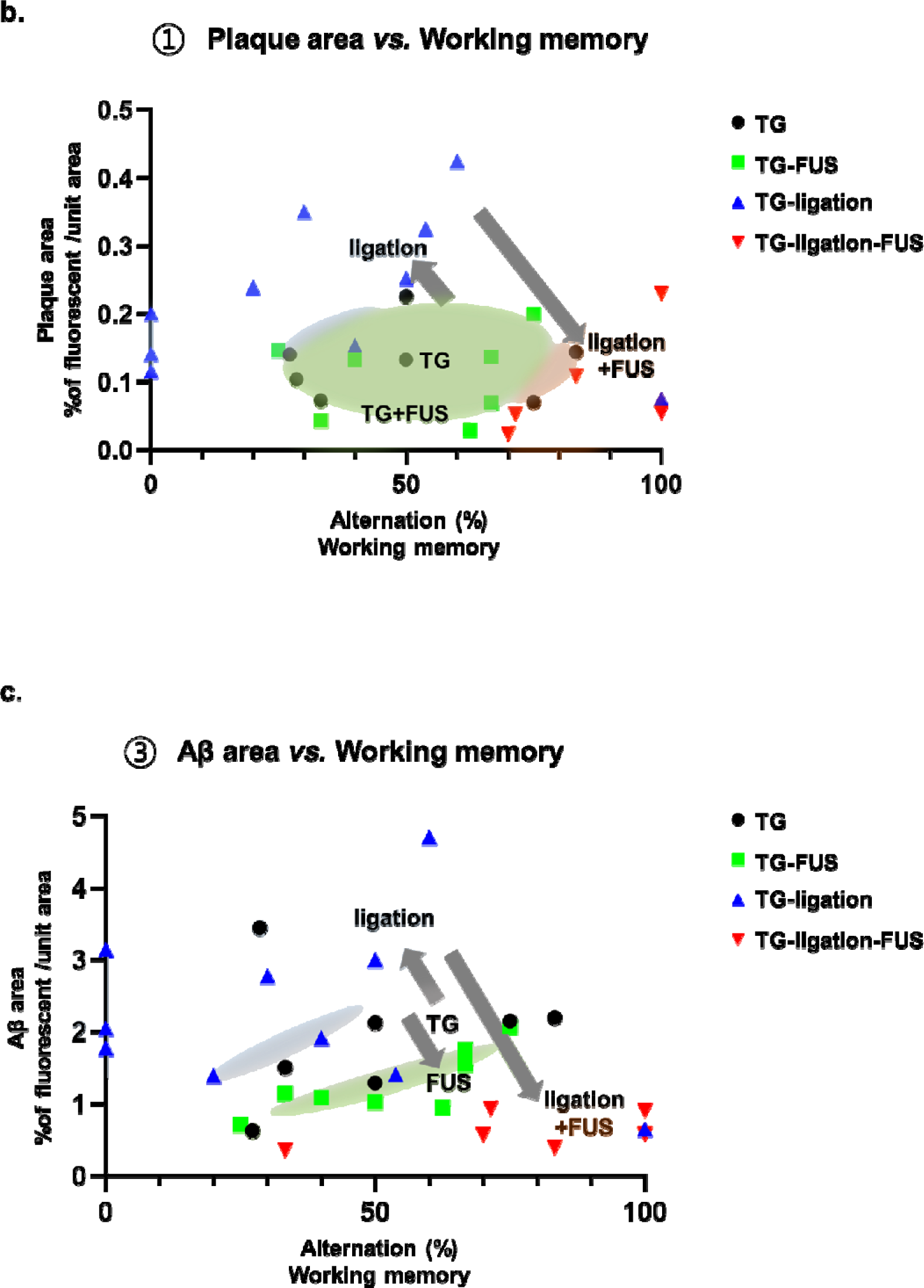
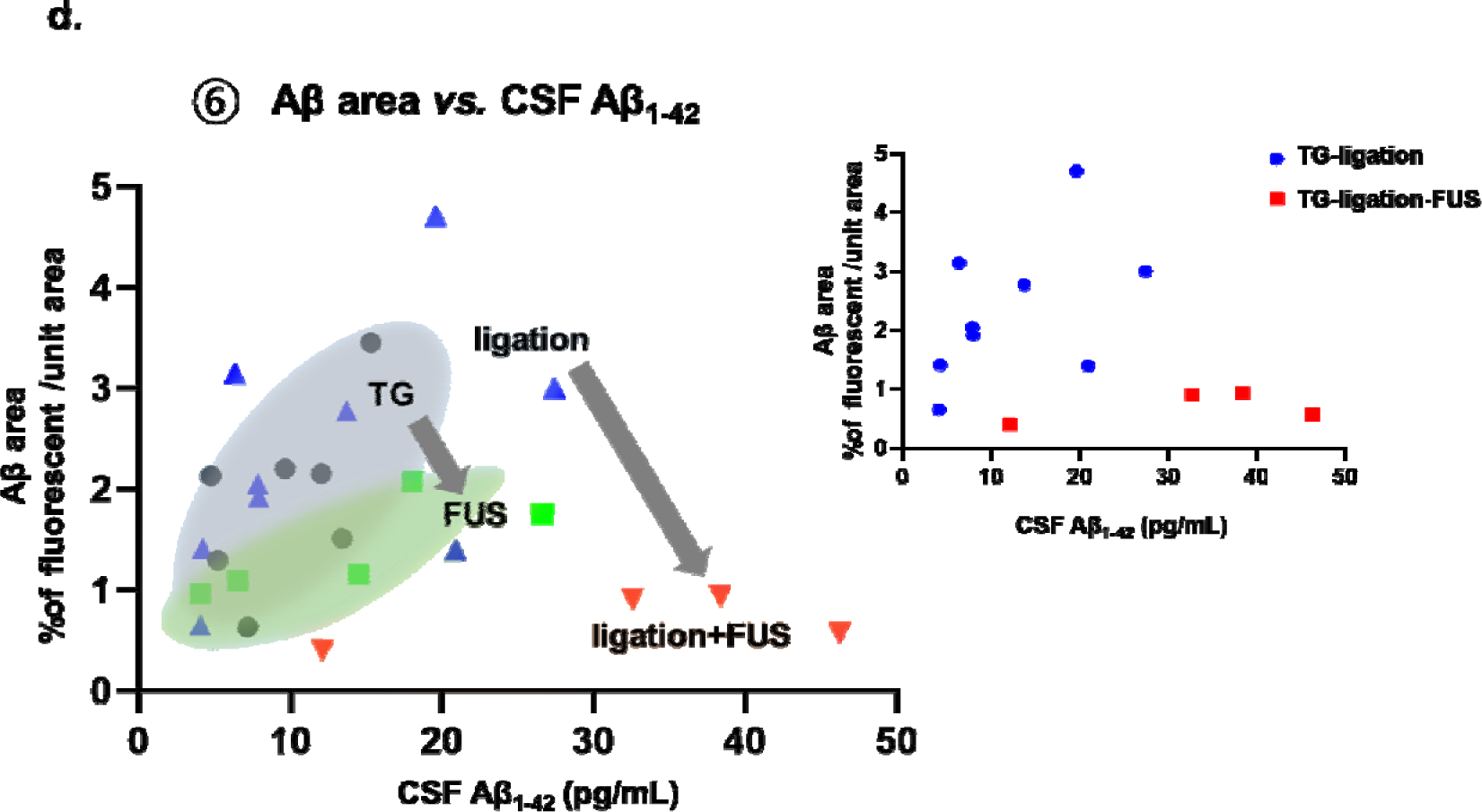
Correlation between measured values from individuals. **a**. The correlation table between measured values from individuals summarized the number of individuals and their *p*-values. *P* values were obtained from simple linear regression between the values (Prism software). Pos Corr and Neg Corr mean positive and negative correlation, respectively. Subgroups include dCLN ligated mice and dCLN ligated mice with FUS-MB. And n.s. means not significant. **b**. Correlation graph between plaque area and working memory (Y-maze) in all individuals. **c**. Correlation graph between Aβ area and working memory in all individuals. **d**. Correlation graph between Aβ area and CSF Aβ in all individuals. Ellipses represent the distribution of individuals in each group. Gray arrows indicate the direction of changes in the group distribution depending on the conditions.

## Materials and Methods

### Animal models

The 5XFAD mice contain five FAD mutations in human APP and PS1, and age matched littermates were used as wild-type controls. One group was set up at the 4 months of age and FUS-MB were treated once a week for 6 weeks. After one week passed after the last FUS-MB, the animals were put to behavior test, sampling of CSF, sacrificed and brain specimen were harvested for pathological analysis. The other experimental groups received bilateral lymphatics ligation to dCLN at the age of 4 months. After two weeks of postoperative stabilization, FUS-MB was treated once a week for 4 weeks. Again after one week, behavior test, sampling of CSF for and pathological analysis was performed including Aβ ELISA and Thioflavin S and immunostaining. In this study, all experiments were conducted with approval Institutional Animal Care and Use Committee at Seoul National University.

### FUS-MB procedure

A total of 100 μl MB (SonoVue^®^, Bracco Suisse S.A., #701299) was infused through the tail vein during the sonication. At the center frequency of 715 kHz, the sonication was 60 s in duration and consisted of 20 ms bursts at a pulse repetition frequency of 1 Hz (2% duty cycle). Acoustic pressure at the targeted area was 0.42 MPa. Target region of sonication was determined by our house-made stereotaxic frame. After the sonication, 2% Evans blue at 200μl was intravenously administered to confirm the BBB opening.

### dCLN ligation model

Mice were anesthetized with 2% isoflurane at 1L/min oxygen flow. And then, the incision site was shaved and sterilized. After longitudinal incision of the neck from the mandible to the sternum, the muscles and fascia were separated from the carotid artery under stereomicroscope. dCLN, located around the carotid artery, were carefully dissected from surrounding tissues and ligated with 9-0 nylon suture. After surgery, the animals were stabilized for two weeks. To confirm the ligation of lymphatics, 5 μl of 2% Evans blue was injected into cisterna magna of mice. Without the ligation of dCLN, CSF and dCLN were stained blue 30 min after the Evans blue injection. But, after the ligation, dCLN was not stained blue (Supplementary Fig. 1).

### Y-maze test for memory

The Y-maze test was carried out one week after the last FUS-MB treatment to evaluate short-term working memory. Each mouse was placed at the center of Y-maze and allowed to move for 8 minutes. Percentage alternation is the number of entries into three arms divided by the maximum possible alternations.

### Enzyme-linked immunosorbent assay (ELISA) for Aβ_1-42_

Mice were anesthetized with 2% isoflurane and 5μ of CSF was collected from CSF in cisterna magna. was measured with Human Aβ Ultrasensitive ELISA Kit (Invitrogen, #KHB3544).

### Immunohistochemistry for Aβ and glia cells

Mice were perfused blood flow with cold 1X phosphate-buffered saline (PBS) and the whole brains were harvested with the skull attached. After carefully separating brains from the skull, the brains were coronally sectioned with 2 mm, embedded in paraffin embedded, and analyzed with 4 μm sections. The 4 μm sectioned brain samples were deparaffinized in xylene and rehydrated in a series of graded ethanol solutions. After antigen retrieval using 0.01M of citric acid for 10 minutes, increase the antibody permeability with 0.5% TritonX-100 in 2.5X tris-buffer saline (TBS) and block the non-specific antigens with 5% bovine serum albumin (BSA). And then brain sections were incubated overnight with the following primary antibodies directed against Aβ (Cell signaling, #8243S), Aβ-6E10 (BioLegend, #803001), GFAP (Cell signaling, #3670S), Iba1 (Abcam, #ab153696), αSMA (Abcam, #ab7817). After overnight incubation, the brain sections were washed and incubated 1 hour with following secondary antibodies (Alexa Fluor 488 Goat anti-mouse IgG, Life technologies, #A11001/ Alexa Fluor 555 donkey anti-rabbit IgG, Life technologies, #A31572). Fully washed sections were mounted. For the capture of stained slide, the LEICA confocal microscopy SP8 was used.

### Thioflavin S staining for amyloid plaques

The paraffin embedded sections were deparaffinized in xylene and rehydrated in a series of graded ethanol solution. The hydrated brain sections were incubated in 1% filtered Thioflavin S solution for 8 minutes and washed with 70% ethanol and distilled water (DW). To quantify the Thioflavin S stained slice images, Image J software was used.

### Hematoxylin and Eosin staining for brain

After deparaffinization and rehydration, sectioned brain tissues were stained with Harris hematoxylin for 30 seconds and washed with DW for 10 minutes. Next, the tissues were stained with eosin for 1 min, dehydrated and washed with xylene. Canada balsam was used for mounting.

### TUNEL staining for brain

To detect apoptosis in brain tissue, DeadEnd^TM^ Fluorometric TUNEL System (Promega, #G3250) was used. The above indexes were quantified according to the manufacturer’s instructions. Briefly, the deparaffinized tissues were fixed and washed. And the tissues were incubated in 20ug/ml proteinase K to increase permeability. Next, the washed tissues were incubated in equilibration buffer and labelled with TdT reaction mixture at 37°C for 60 minutes. After the reaction, the slides were immersed in 2X SSC to stop the reaction and washed.

### Statistical analysis

Data were statistically analyzed with Prism and Real Statistics/Excel softwares. Values were reported as average ± standard deviation (SD). One-way ANOVA test with post hoc Tukey was used for multiple comparisons. And unpaired t-test was used for comparison of wild type and 5XFAD groups in Y-maze test. Significant difference was assigned as *p* < 0.05 (*), *p* < 0.005 (**).

